# ReConPlot – an R package for the visualization and interpretation of genomic rearrangements

**DOI:** 10.1101/2023.02.24.529890

**Authors:** Jose Espejo Valle-Inclán, Isidro Cortés-Ciriano

## Abstract

Whole-genome sequencing studies of human tumours have revealed that complex forms of structural variation, collectively known as complex genome rearrangements (CGR), are pervasive across diverse cancer types. Detection, classification, and mechanistic interpretation of CGR requires the visualization of complex patterns of somatic copy number aberrations (SCNAs) and structural variants (SVs). However, there is a lack of tools specifically designed to facilitate the visualization and study of CGR. We present **ReConPlot** (**RE**arrangement and **CO**py **N**umber **PLOT**), an R package that provides functionalities for the joint visualization of SCNAs and SVs across one or multiple chromosomes. ReConPlot is based on the popular ggplot2 package, thus allowing customization of plots and the generation of publication-quality figures with minimal effort. Overall, ReConPlot facilitates the exploration, interpretation, and reporting of complex genome rearrangement patterns.

**Code availability:** The R package ReConPlot is available at https://github.com/cortes-ciriano-lab/ReConPlot. Detailed documentation and a tutorial with examples are provided with the package.

## 1 Introduction

The advent of whole genome sequencing (WGS) has enabled a more nuanced characterization of the diversity, rates and underlying mechanisms of chromosomal alterations than was ever possible using cytogenetic or pathology analyses^1–4^. WGS studies of human cancers have revealed that genomic instability, a hallmark of cancer, manifests as alterations in the structure and number of chromosomes (aneuploidy), whole genome doubling (WGD), repeat instability, and remarkably diverse forms of structural variants (SVs)^5–8^. SVs, which account for most driver events in some cancer types^5,9^, refer to the rearrangement of the genome leading to the deletion, amplification or reshuffling of genomic segments. In cancer genomes, genomic rearrangements manifest as (1) simple events, such as deletions, duplications, inversions, and insertions occurring in isolation, or (2) complex events involving multiple breakpoints across one or multiple chromosomes and showing complex patterns of both spatial and temporal clustering^10–13^. Such complex patterns, collectively referred to as complex genomic rearrangements (CGR), include those recently discovered in cancer genome studies, such as chromothripsis^11,14,15^, chromoanasynthesis^16^, chromoplexy^17^, pyrgo, rigma, and tyfonas^13^, as well as others initially described in cytogenetic studies, such as breakage–fusion–bridge (BFB) cycles^18^ and extrachromosomal DNA elements^19,20^. Multiple algorithms have been developed to detect and classify CGR^11–13,21^ based on the analysis of the patterns of SVs and SCNAs detected through computational cancer genome analysis. However, due to the diversity, complexity, variable scale and overlapping features of CGR, coupled to their co-localization^11^, their detection and classification remains a challenging task. As a result, manual inspection of SCNA and SV data is often required to resolve the most complex cases^11,21,22^. This task requires versatile methods to visualize SCNAs and SVs across genomic regions ranging from a few kbp to multiple chromosomes. A popular approach for genomics data visualization, termed Circos plot^23^, allows exploration of CGR by displaying the cancer genome in a circular layout where concentric circles show different types of mutations and genomic features^24^. Although versatile to provide an overview of the cancer genome^25–27^, Circos plots are often too complex to visualize CGR involving large numbers of SVs and SCNAs. An alternative approach consists of displaying genomic regions of interest in a linear layout where regions of equal copy number are represented by segments, and SVs by arcs^5,22^. This visualization strategy, usually referred to as genomic rearrangement plots, has become popular in the cancer genomics community for visualizing and reporting the patterns and consequences of CGR (*e.g*., disruption of tumour suppressor genes by SVs)^5,11^. However, there is lack of easy-to-use software packages to visualize genomic rearrangement profiles and generate publication-quality figures for reporting cancer genome analysis results. Here we present **ReConPlot** (**RE**arrangement and **CO**py **N**umber **PLOT**), an R package that provides functionalities for the joint visualization of SCNAs and SVs across one or multiple chromosomes.

## 2 Methods

ReConPlot relies on the popular package ggplot2^28^ for the visualization of SCNA and SV profiles, thus allowing for user-specific customization and integration with functionalities from other R packages to compose multi-panel figures easily. The main function of the package, *ReConPlot*, only requires as input the genomic coordinates for the regions to be visualized, integer minor and total copy number data, and SV information in browser extensible data paired-end (BEDPE) format^29^. *ReConPlot* permits the visualization of genomic rearrangement profiles across one or multiple chromosomes (Fig. 1). Each ReConPlot consists of three main panels. The bottom panel shows Giemsa binding data^30,31^ for the genomic regions of interest. The middle panel reports total and minor copy number information. Finally, the top panel shows SVs. SVs whose breakpoints fall within the regions selected to be displayed are represented by arcs. In cases where only one breakpoint maps to the selected genomic regions, the SV is represented as a vertical line ending with a 45-degree overhang. SVs are categorized into four groups depending on the read orientation at the breakpoints (i.e., type of fragment joins) following the notation established by the Pan-Cancer Analysis of Whole Genomes project (PCAWG^5^; Fig. 1): deletion-like SVs (DEL) are represented as “+ −”, duplication-like SVs (DUP) as “− +”, tail-to-tail inversions (t2tInv) as “− −”, and head-to-head inversions (h2hInv) as “+ +”. Using the same notation as PCAWG allows for smooth integration with other software packages designed for the detection, classification and interpretation of CGR, such as ShatterSeek^11^. In addition, *ReConPlot* provides functionalities to highlight the location of genes (see Fig. 1 for examples). Currently, ReConPlot supports the following builds of the human reference genome: GRCh37, GRCh38 and T2T-CHM13. While default parameter values ensure the generation of publication-quality figures, the function *ReConPlot* is highly versatile, as it allows customization of the layout of the plots, font sizes, font colours, and other graphical parameters (see the documentation and tutorial of the package for a full list of customizable graphical options).

**Figure 1.**
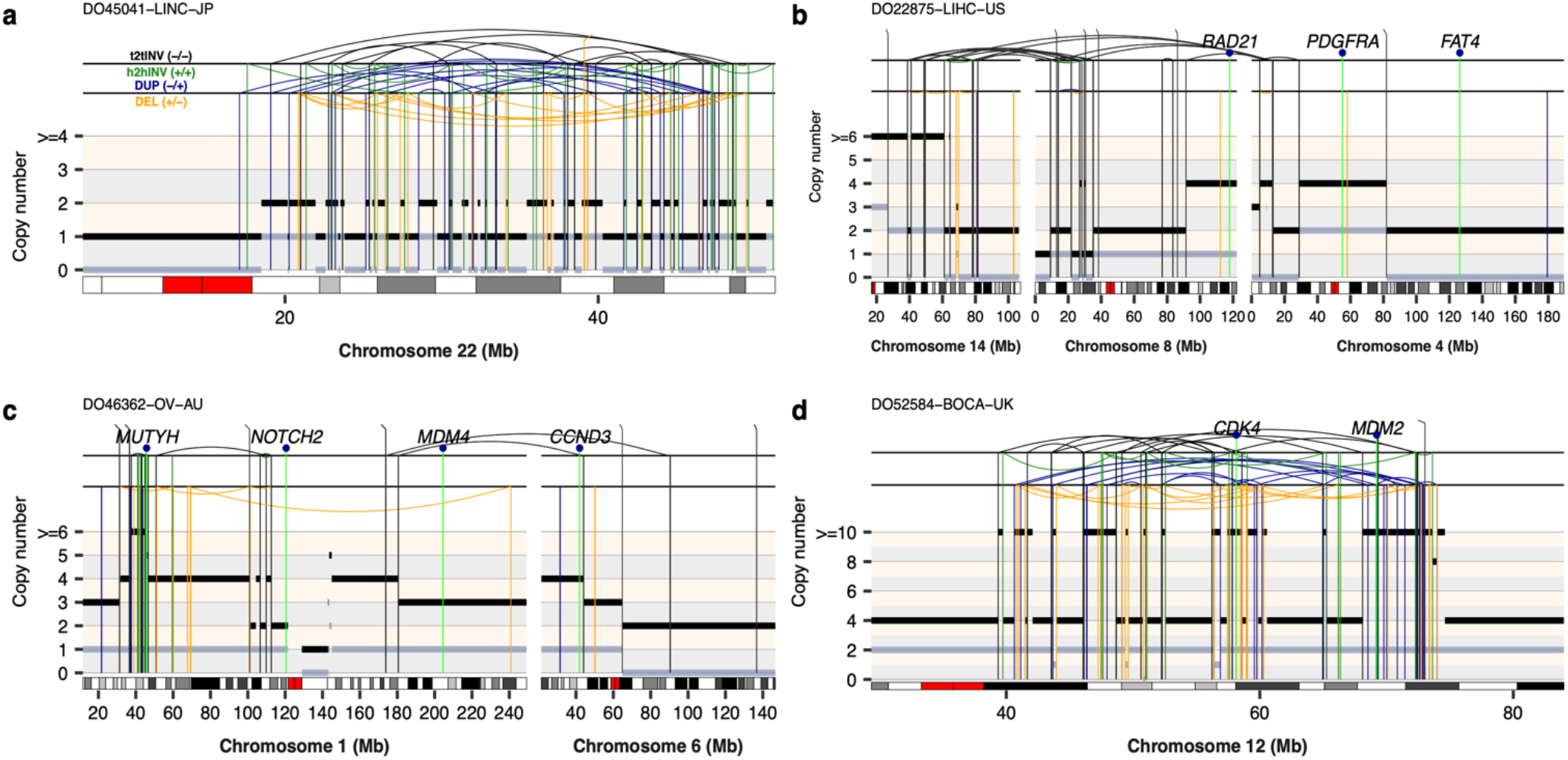
Examples of ReConPlots visualizing complex genomic rearrangements detected in four cancer genomes from the PCAWG cohort. (**a**) Example of a canonical chromothripsis event detected in a liver adenocarcinoma. The ReConPlot shows the characteristic cluster of interleaved SVs and copy number oscillations between two copy number states accompanied by loss of heterozygosity, which is indicated by those regions with a minor copy number of 0. (**b-c**) Examples of CGR spanning multiple chromosomes detected in a liver adenocarcinoma and an ovarian adenocarcinoma, respectively. (**d**) Example of a chro-mothripsis event detected in an osteosarcoma genome. The chromothripsis event occurred after whole-genome doubling, as evidenced by the minor copy number oscillations between 1 and 2, and caused the high-level amplification of *CDK4* and *MDM2*. Tail-to-tail (t2tINV) inversion, head-to-head (h2hINV) inversions, duplication-like SVs (DUP), and deletion-like SVs (DEL) are depicted in black, green, blue, and orange, respectively. Total and minor copy number values are represented by black and grey segments, respectively. ICGC IDs are shown on top of each ReConPlot.

## 3 Results

We have extensively validated the functionalities of ReConPlot using SV and SCNA calls from the PCAWG project, allowing us to identify and classify diverse CGR, such as chromothripsis, CGR involving multiple chromosomes, and CGR showing high-level oncogene amplifications (Fig. 1). In sum, ReConPlot provides functionalities for the visualization and interpretation of complex genomic rearrangement profiles detected in cancer genomes and rare diseases patients.

## Acknowledgements

J.E.V.-I. and I.C.-C. thank the European Molecular Biology Laboratory (EMBL) for funding

## Conflict of Interest

The authors declare no conflicts of interest.

## Notes

https://github.com/cortes-ciriano-lab/ReConPlot

### Competing Interest Statement

The authors have declared no competing interest.

### Summary of Updates

Fixed GitHub link.

## References

1. Mardis, E. R. & Wilson, R. K. Cancer genome sequencing: a review. Hum. Mol. Genet. 18, R163–8 (2009).

2. Greenman, C. et al. Patterns of somatic mutation in human cancer genomes. Nature 446, 153–158 (2007).

3. Garraway, L. A. & Lander, E. S. Lessons from the cancer genome. Cell 153, 17–37 (2013).

4. Cortés-Ciriano, I., Gulhan, D. C., Lee, J. J.-K., Melloni, G. E. M. & Park, P. J. Computational analysis of cancer genome sequencing data. Nat. Rev. Genet. (2021) doi:10.1038/s41576-021-00431-y.

5. ICGC/TCGA Pan-Cancer Analysis of Whole Genomes Consortium. Pan-cancer analysis of whole genomes. Nature 578, 82–93 (2020).

6. Priestley, P. et al. Pan-cancer whole-genome analyses of metastatic solid tumours. Nature 575, 210–216 (2019).

7. Steele, C. D. et al. Signatures of copy number alterations in human cancer. Nature 606, 984–991 (2022).

8. Macintyre, G. et al. Copy number signatures and mutational processes in ovarian carcinoma. Nat. Genet. 50, 1262–1270 (2018).

9. Zack, T. I. et al. Pan-cancer patterns of somatic copy number alteration. Nat. Genet. 45, 1134–1140 (2013).

10. Li, Y. et al. Patterns of somatic structural variation in human cancer genomes. Nature 578, 112–121 (2020).

11. Cortés-Ciriano, I. et al. Comprehensive analysis of chromothripsis in 2,658 human cancers using whole-genome sequencing. Nat. Genet. 52, 331–341 (2020).

12. Bao, L., Zhong, X., Yang, Y. & Yang, L. Starfish infers signatures of complex genomic rearrangements across human cancers. Nat. Cancer 3, 1247–1259 (2022).

13. Hadi, K. et al. Distinct classes of complex structural variation uncovered across thousands of cancer genome graphs. Cell 183, 197–210.e32 (2020).

14. Stephens, P. J. et al. Massive genomic rearrangement acquired in a single catastrophic event during cancer development. Cell 144, 27–40 (2011).

15. Rausch, T. et al. Genome sequencing of pediatric medulloblastoma links catastrophic DNA rearrangements with TP53 mutations. Cell 148, 59–71 (2012).

16. Liu, P. et al. Chromosome catastrophes involve replication mechanisms generating complex genomic rearrangements. Cell 146, 889–903 (2011).

17. Baca, S. C. et al. Punctuated evolution of prostate cancer genomes. Cell 153, 666–677 (2013).

18. Campbell, P. J. et al. The patterns and dynamics of genomic instability in metastatic pancreatic cancer. Nature 467, 1109–1113 (2010).

19. Deshpande, V. et al. Exploring the landscape of focal amplifications in cancer using AmpliconArchitect. Nat. Commun. 10, 392 (2019).

20. Turner, K. M. et al. Extrachromosomal oncogene amplification drives tumour evolution and genetic heterogeneity. Nature 543, 122–125 (2017).

21. Notta, F. et al. A renewed model of pancreatic cancer evolution based on genomic rearrangement patterns. Nature 538, 378–382 (2016).

22. Li, Y. et al. Constitutional and somatic rearrangement of chromosome 21 in acute lymphoblastic leukaemia. Nature 508, 98–102 (2014).

23. Krzywinski, M. et al. Circos: an information aesthetic for comparative genomics. Genome Res. 19, 1639–1645 (2009).

24. Nusrat, S., Harbig, T. & Gehlenborg, N. Tasks, techniques, and tools for genomic data visualization. Comput. Graph. Forum 38, 781–805 (2019).

25. Davies, H. et al. HRDetect is a predictor of BRCA1 and BRCA2 deficiency based on mutational signatures. Nat. Med. 23, 517–525 (2017).

26. Goldman, M. et al. A user’s guide to the online resources for data exploration, visualization, and discovery for the Pan-Cancer Analysis of Whole Genomes project (PCAWG). bioRxiv (2017) doi:10.1101/163907.

27. Shale, C. et al. Unscrambling cancer genomes via integrated analysis of structural variation and copy number. Cell Genom. 2, 100112 (2022).

28. Wickham, H. ggplot2. (Springer International Publishing, 2016).

29. Quinlan, A. R. & Hall, I. M. BEDTools: a flexible suite of utilities for comparing genomic features. Bioinformatics 26, 841–842 (2010).

30. Furey, T. S. & Haussler, D. Integration of the cytogenetic map with the draft human genome sequence. Hum. Mol. Genet. 12, 1037–1044 (2003).

31. Cheung, V. G. et al. Integration of cytogenetic landmarks into the draft sequence of the human genome. Nature 409, 953–958 (2001).

